# Sequentially activated death complexes regulate pyroptosis and IL-1β release in response to *Yersinia* blockade of immune signaling

**DOI:** 10.1101/2023.09.14.557714

**Authors:** Ronit Schwartz Wertman, Christina K. Go, Benedikt S. Saller, Olaf Groß, Phillip Scott, Igor E. Brodsky

## Abstract

The *Yersinia* virulence factor YopJ potently inhibits immune signaling in macrophages by blocking activation of the signaling kinases TAK1 and IKK. In response, macrophages trigger a backup pathway of host defense that mediates cell death via the apoptotic enzyme caspase-8 and pyroptotic enzyme caspase-1. While caspase-1 is normally activated within multiprotein inflammasome complexes that contain the adaptor ASC and NLRs, which act as sensors of pathogen virulence, caspase-1 activation following *Yersinia* blockade of TAK1/IKK surprisingly requires caspase-8 and is independent of all known inflammasome components. Here, we report that caspase-1 activation by caspase-8 requires both caspase-8 catalytic and auto-processing activity. Intriguingly, while caspase-8 serves as an essential initiator of caspase-1 activation, caspase-1 amplifies its own activation through a feed-forward loop involving auto-processing, caspase-1-dependent cleavage of the pore-forming protein GSDMD, and subsequent activation of the canonical NLRP3 inflammasome. Notably, while caspase-1 activation and cell death are independent of inflammasomes during *Yersinia* infection, IL-1β release requires the canonical NLPR3 inflammasome. Critically, activation of caspase-8 and activation of the canonical inflammasome are kinetically and spatially separable events, as rapid capase-8 activation occurs within multiple foci throughout the cell, followed by delayed subsequent assembly of a single canonical inflammasome. Importantly, caspase-8 auto-processing normally serves to prevent RIPK3/MLKL-mediated necroptosis, and in caspase-8’s absence, MLKL triggers NLPR3 inflammasome activation and IL-1β release. Altogether, our findings reveal that functionally interconnected but temporally and spatially distinct death complexes differentially mediate pyroptosis and IL-1β release to ensure robust host defense against pathogen blockade of TAK1 and IKK.

**One Sentence Summary:** *Yersinia*-induced cell death and IL-1β release are driven by spatially and temporally distinct but functionally connected death complexes.

## INTRODUCTION

The innate immune system is critical for host defense against bacterial pathogens, as it detects pathogen-associated molecular patterns (PAMPs) as well as pathogen-mediated perturbations of host biological pathways^1,2^. Apoptosis, pyroptosis and necroptosis are distinct forms of regulated cell death that mediate anti-microbial host defense^3,4^. Apoptosis is classically viewed as a developmentally programmed or homeostatic, non-inflammatory cell death, whereas pyroptosis is a lytic form of cell death accompanied by release of inflammatory IL-1 family cytokines that takes place in response to microbial infection^5–7^. Apoptosis and pyroptosis are both driven through activation of caspases, pro-enzyme cysteine proteases that undergo proteolytic activation following recruitment to multiprotein complexes^8^, while necroptosis is caspase-independent^9,10^.

Apoptosis and pyroptosis require engagement of distinct signaling complexes and effector caspases, and are traditionally thought to be mutually exclusive and cross-inhibitory^3^. However, disruption of core immune signaling pathways by pathogen virulence factors can trigger cell death that exhibits biochemical features of both apoptosis and pyroptosis^11^. Indeed, recent studies have proposed the existence of a cell death pathway involving simultaneous activation of pyroptosis, apoptosis, and necroptosis, termed PANoptosis^12,13^ following microbial infection or disruption of immune signaling pathways. However, as the morphologic and physiologic consequences of distinct cell death pathways are unique, and the effector enzymes of one death pathway typically cross-inhibit the others, how an individual cell might simultaneously undergo multiple distinct forms of cell death is unclear.

During apoptosis, processing of executioner caspases-3 and −7 by the initiator caspase-8 results in the cleavage of numerous caspase-3/7-dependent substrates, leading to the organized breakdown of the cell into membrane-enclosed ‘blebs’ that are rapidly phagocytosed by neighboring cells with minimal inflammation^5^. Conversely, during pyroptosis, caspase-1 is activated by its recruitment into inflammasomes, multiprotein signaling complexes that form in response to microbial contamination of the cytosol^6,7^, and are nucleated by sensor NLR proteins and the adaptor protein ASC (apoptosis-associated speck like protein containing a caspase activation and recruitment domain)^14^. Active caspase-1 processes the inflammatory cytokine pro-IL-1β and pore-forming protein Gasdermin D (GSDMD). The N-terminal fragment of GSDMD (p30) oligomerizes and inserts into the plasma membrane, releasing mature IL-1β as well as other intracellular alarmins through membrane rupture and cell lysis^14–16^. However, pathogenic *Yersiniae* inject a variety of virulence factors known as *Yersinia* outer proteins (Yops) into the cytoplasm of host cells through their Type 3 Secretion Systems (T3SS)^17–19^ to disrupt innate immune responses. Among these is the acetyl-transferase YopJ, which blocks IKK and TAK1 signaling^20–22^. Such blockade leads to the combined activation of caspase-1 and caspase-8, and elicits caspase-1 and caspase-8-dependent cleavage of GSDMD and IL-1β^23,24^. Interestingly, caspase-1 activation following *Yersinia pseudotuberculosis* (*Yptb*) infection is independent of all currently known inflammasome components, including NLRP3, NLRC4 and the inflammasome adaptor protein ASC^25^, but is instead dependent on caspase-8^25^. Intriguingly, despite the lack of a requirement for ASC in caspase-8 or −1 activation or cell death^25^, ASC forms large oligomers in response to YopJ activity, suggesting that ASC complexes play an as-yet-undefined role in *Yersinia* infection^24^. While the ASC pyrin (PYD) domain interacts with the caspase-8 death-effector (DED) domain^24,26,27^, whether these distinct pathways are activated simultaneously or sequentially within infected cells and their role in promoting programmed cell death and inflammatory responses is poorly defined.

Here, we find that caspase-8-dependent caspase-1 activation requires both caspase-8 and caspase-1 activity. Surprisingly, despite the ability of caspase-8 to cleave caspase-1 directly, caspase-1 catalytic activity was required for its own processing downstream of caspase-8 activation, indicating that caspase-1 acts as a feed-forward amplifier of caspase-8-dependent pyroptosis. Macrophages that express an uncleavable caspase-8 (*Casp8^D387A/D387A^)* are sensitized to RIPK3-mediated necroptosis, which triggers a backup pathway of caspase-1 activation to enable pyroptotic cell death and IL-1β release even in the absence of active caspase-8. Additionally, although ASC is not required for caspase-1 activation during *Yptb* infection, IL-1β release requires the canonical NLRP3 inflammasome. These findings indicate that secondary NLRP3 inflammasome activation subsequent to GSDMD cleavage and potassium-efflux mediates IL-1β release. Indeed, caspase-8 activation preceded assembly of ASC puncta, and ASC puncta and active caspase-8 were differentially localized within macrophages. Altogether, this work demonstrates that functionally linked, but temporally and spatially distinct death complexes mediate pyroptosis and IL-1β release in response to pathogen blockade of innate immune signaling.

## RESULTS

### Caspase-8 activity is required for cell death and caspase-1 processing

During *Yersinia pseudotuberculosis* (*Yptb*) infection, cell death and caspase-1 processing occur independently of all known inflammasome components^25^. Consistent with previous findings from our group and others^25,28^, cell death in response to *Yersinia* YopJ activity is dependent on caspase-8, as in contrast to either *Ripk3^−/−^* or C57BL/6 bone marrow-derived macrophages (BMDMs), *Casp8^−/−^Ripk3^−/−^* BMDMs remain viable following *Yptb* infection **(FIG. 1A)**. Furthermore, processing of caspase-1 into its active p20 fragment is dependent on caspase-8, as in contrast to *Ripk3^−/−^* BMDMs, it is not observed in *Casp8^−/−^Ripk3^−/−^* BMDMs, **(FIG. 1B)**. Consistent with prior findings^23,24,29^, GSDMD processing also requires caspase-8 but not RIPK3 (**FIG. 1B**). To determine if caspase-8 is sufficient for caspase-1 activation, as well as to define its molecular requirements, we co-expressed caspase-1 with various caspase-8 constructs in which the caspase-8 death-effector domains (DEDs) were replaced with an inducible dimerizable domain that promotes its activation upon addition of the dimerizer AP20187^30^ **(FIG. 1C)**. Addition of AP20187 to induce dimerization of caspase-8 triggered robust caspase-1 processing into its active p20 fragment, which was undetectable in the absence of dimerizer **(FIG. 1D)**. Critically, dimerizable constructs containing catalytic mutant caspase-8 (C360A) or uncleavable caspase-8 lacking five aspartate processing sites^30^ were unable to promote caspase-1 cleavage, indicating that both caspase-8 catalytic activity and auto-processing are required for caspase-1 cleavage **(FIG. 1D)**. To determine if auto-processed caspase-8 functions solely as a scaffold to recruit caspase-1, or whether its catalytic activity is necessary for caspase-1 processing, we expressed dimerizable caspase-8 constructs in which the interdomain auto-processing site at position D384 was replaced with the cleavage sequence for the tobacco etch virus (TEV) protease. While addition of dimerizer to cells co-transfected with caspase-8-TEV and caspase-1 led to some baseline caspase-1 processing, co-expression of TEV protease to allow for caspase-8 cleavage resulted in maximal caspase-1 processing **(FIG. 1E)**. Notably, caspase-8 catalytic activity was essential for caspase-1 activation even when caspase-8 was dimerized and exogenously cleaved by TEV, demonstrating that both caspase-8 cleavage and enzymatic activity are absolutely required for caspase-1 processing **(FIG. 1E)**.

**Figure 1:**
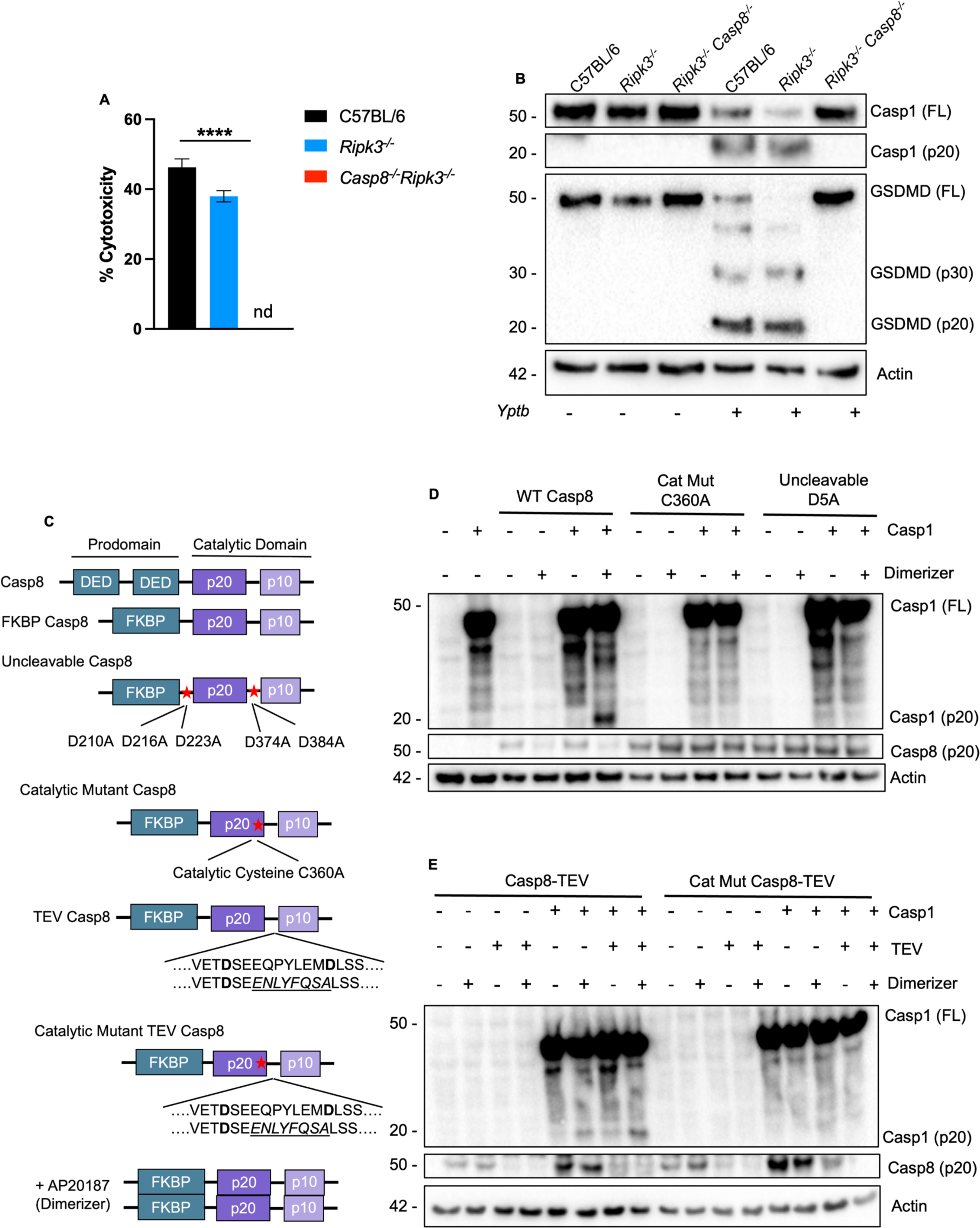
Caspase-8 activity is required for cell death and caspase-1 processing. (A) C57BL/6, *Ripk3^−/−^*, and *Casp8^−/−^Ripk3^−/−^* BMDMs were infected with WT *Yptb* and percent cytotoxicity was measured 4 hours post-infection as described in materials and methods. (B) Lysates collected 3 hours post-infection were immunoblotted for caspase-1 and GSDMD. (C) Schematic representation of FKBP constructs of caspase-8 employed in this study. (D) HEK293T cells transfected with caspase-1 and WT, catalytically inactive (C360A), or uncleavable (D5A) FKBP-caspase-8 and induced to dimerize with AP20187 (dimerizer) 24 hours post transfection. Lysates were collected for western blotting 6 hours after adding AP20187. (E) HEK293T cells transfected with caspase-1, TEV, and WT or catalytically inactive (C360A) FKBP-caspase-8-TEV and treated with dimerizer as indicated 24 hours post transfection. Lysates were collected for western blotting 6 hours after AP20187 addition. Nd — not detected, **** p < 0.0001 by two-way ANOVA. Error bars represent the mean +/-SEM of triplicate wells and are representative of three independent experiments.

### Caspase-1 activation by caspase-8 requires caspase-1 catalytic activity

Our findings indicate that caspase-8 acts as an apical initiator caspase to activate caspase-1 in response to blockade of TAK1 and IKK signaling by pathogens. Caspase-1 autoproteolysis is required for its activation within canonical inflammasomes^31–33^. In contrast, during apoptosis, caspase-8 processes caspase-3 into its mature form, but caspase-3 does not undergo autoproteolysis, thus limiting its feed-forward amplification capacity^33,34^. Unexpectedly, however, catalytically inactive caspase-1 (Casp1^C284A^) failed to undergo processing in response to inducible dimerization of caspase-8 in HEK293T cells, indicating that caspase-1 catalytic activity was required for its own processing in the setting of caspase-8 activation **(FIG. 2A).** Notably, while *Casp1^−/−^* immortalized BMDMs (iBMDMs) stably expressing WT caspase-1 robustly processed caspase-1 upon infection with *Salmonella* Typhimurium or *Yptb* **(FIG. 2B)**, iBMDMs expressing catalytically inactive caspase-1 DEAD (C284A)^32^ were unable to process caspase-1 during either *Salmonella* or *Yptb* infection. These observations support our findings that caspase-1 catalytic activity is necessary for its own processing and activation downstream of YopJ-induced caspase-8 activation **(FIG. 2B)**. Consistent with previous findings that *Yptb-*induced cell death does not require caspases-1 or −11^24,25^, caspase-1 DEAD cells exhibited wild-type levels of LDH release upon infection with *Yptb* **(FIG. 2C)**, but failed to induce LDH release upon infection with *S.* Typhimurium, as expected **(FIG. 2C, S1A)**. Critically, primary BMDMs from knock-in mice lacking caspase-1 catalytic activity (*Casp1^mlt/mlt^*)^35^ also failed to process caspase-1, and had significantly reduced processing of GSDMD, as well as reduced IL-β processing and release in response to *Yptb* infection (**FIG. 2D-F**). Similarly to iBMDMs expressing catalytically inactive caspase-1 DEAD (C284A) however, cytotoxicity responses to *Yptb* infection were normal, despite being unable to undergo cytotoxicity in response to *Salmonella* (**FIG. 2G, S1B**). In contrast to IL-1β release, IL-12 secretion by *Casp1^mlt/mlt^* BMDMs was largely intact **(FIG. S1C**). Taken together, our results show that caspase-8 and caspase-1 enzymatic activities are both critical for caspase-1 processing in response to *Yptb* infection, and that caspase-1 catalytic activity is required for IL-1β secretion, even in the presence of sufficient caspase-8 activity.

**Figure 2:**
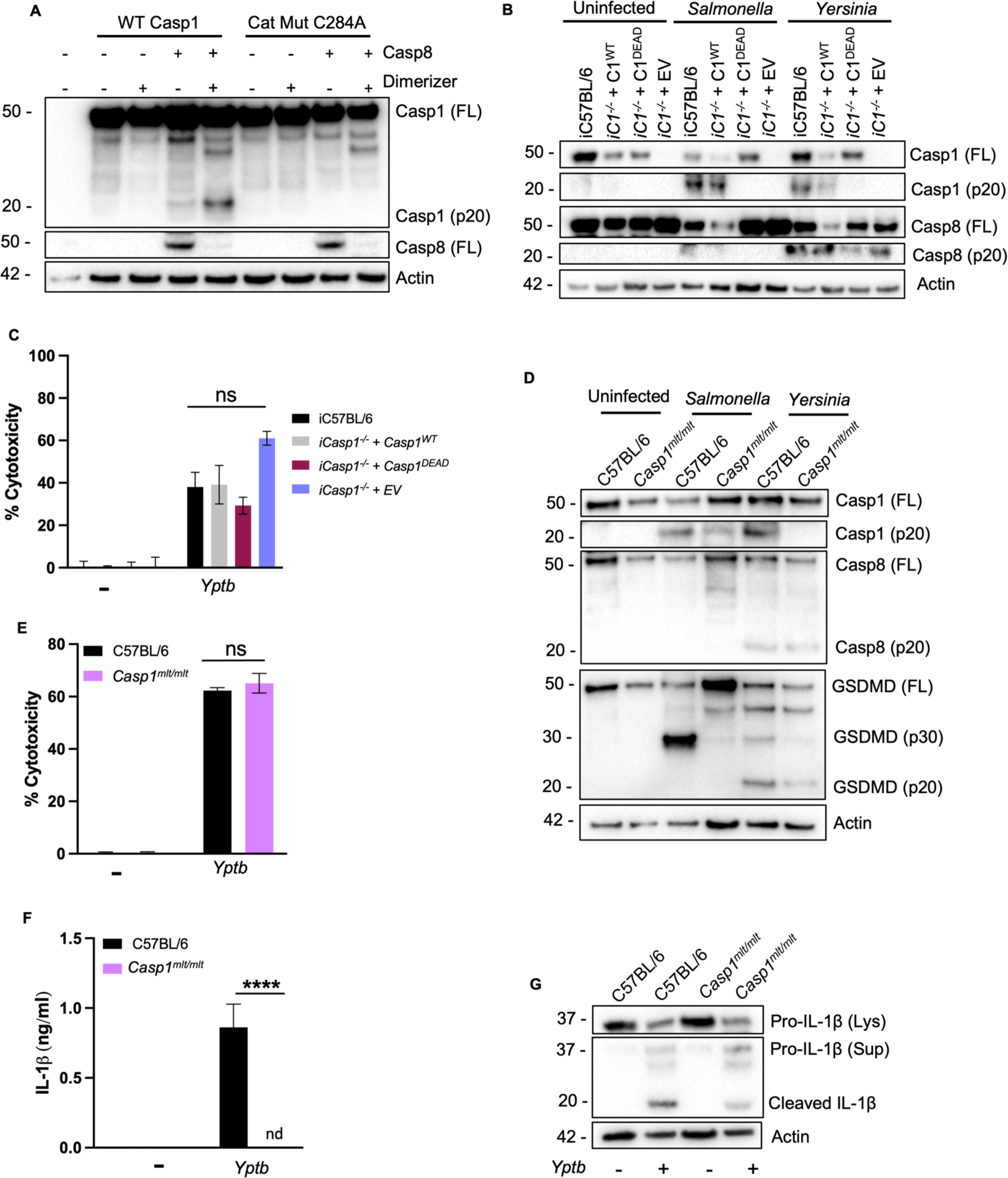
Caspase-1 activation by caspase-8 requires caspase-1 catalytic activity. (A) HEK293T cells were transfected with FKBP-caspase-8 and WT or catalytically inactive (C284A) caspase-1 and treated with dimerizer. Lysates were collected for western blotting as described in materials and methods. (B) iC57BL/6, *iCasp1^−/−^ + Casp1^WT^*, *iCasp1^−/−^ + Casp1^DEAD^,* and *iCasp1^−/−^ + EV* immortalized BMDMs were infected with WT *Yptb* as described in materials and methods. Lysates collected 3 hours post-infection were immunoblotted for caspase-1, caspase-8, GSDMD, and β-actin as indicated. (C) Percent cytotoxicity was assayed 4 hours post-infection as described in materials and methods. (D) C57BL/6, *Casp1^mlt/mlt^* BMDMs were infected with WT *Yptb* as described. Lysates collected 3 hours post-infection were immunoblotted for caspase-1, caspase-8, GSDMD, and β-actin. (E) Percent cytotoxicity was measured 4 hours post-infection. (F) Release of IL-1β into the supernatant was measured by ELISA 4 hours post-infection. (G) Lysates and supernatants collected 3 hours post-infection were immunoblotted for IL-1β. ns — not significant, **** p < 0.0001 by two-way ANOVA. Error bars represent the mean +/-SEM of triplicate wells and are representative of three independent experiments.

### Caspase-8 auto-processing limits RIPK3-mediated necroptosis

*Yersinia* infection or TAK1 blockade have been proposed to induce a combined form of cell death termed PANoptosis, involving the simultaneous activation of pyroptosis, apoptosis, and necroptosis, as assessed by phosphorylation of RIPK3 and the Mixed Lineage Kinase Domain Like Pseudokinase (MLKL) pore-forming protein, coincident with activation of apoptotic and pyroptotic caspases^12,13,36^. However, in the absence of caspase-8 auto-processing, cells undergo RIPK3-dependent necroptosis mediated by RIPK3-dependent activation of MLKL^37,38^. Because MLKL pore formation can promote potassium efflux, a common trigger of the NLRP3 inflammasome^37,39,40^, we hypothesized that caspase-1 activation in the absence of caspase-8 auto-processing could result from NLRP3 activation downstream of RIPK3-and MLKL-mediated necroptosis. We therefore monitored cell death and caspase-1 processing in *Casp8^D387A/D387A^* BMDMs, which express an uncleavable caspase-8, either after infection with *Yptb* or treatment with LPS/IKK inhibitor (IKKi), which pharmacologically mimics the activity of YopJ^41^. In contrast to the HEK293T system or in our previous studies in which non-cleavable caspase-8 was expressed in cells lacking RIPK3^25^, we found that *Casp8^D387A/D387A^* BMDMs infected with *Yptb* or treated with LPS/IKKi exhibited comparable LDH release and caspase-1 processing as WT BMDMs (**FIG. 3A-C).** Moreover, *Casp8^D387A/D387A^* only processed GSDMD into the active p30 fragment, whereas WT BMDMs processed GSDMD into both p30 and p20 fragments (**FIG. 3C**). The GSDMD p20 fragment is generated by caspase-3-mediated cleavage^42^, indicating that both caspase-1 and caspase-3 are active in WT macrophages, but only caspase-1 is active in *Casp8^D387A/D387A^*macrophages. Moreover, the RIPK3 inhibitor GSK’872 inhibited both cell lysis and caspase-1 processing in *Casp8^D387A/D387A^* but not WT BMDMs following *Yptb* infection or LPS/IKKi treatment, suggesting that caspase-8 auto-processing during *Yersinia* infection or IKK/TAK1 blockade normally limits RIPK3-mediated necroptosis and subsequent activation of caspase-1 (**FIG. 3A-C**). Notably, while the NLRP3-specific inhibitor MCC950 did not inhibit cell lysis in the *Casp8^D387A/D387A^* BMDMs, it completely blocked caspase-1 processing in *Yptb-*infected or LPS/IKKi-treated *Casp8^D387A/D387A^* BMDMs, indicating that caspase-1 processing downstream of RIPK3 activation is mediated by NLRP3 (**FIG. 3D-F**). Importantly, *Casp8^D387A/D387A^Mlkl^−/−^* BMDMs^43^ exhibited neither cell lysis, caspase-1, caspase-8, nor GSDMD processing, indicating that MLKL activation occurs upstream of caspase-1 activation, GSDMD processing, and cell lysis in *Casp8^D387A/D387A^* BMDMs (**FIG 3A-F**). Notably, we did not observe RIPK3 or MLKL phosphorylation in WT BMDMs following *Yptb* infection **(FIG. 3G**), consistent with the lack of requirement for RIPK3 in *Yptb-*induced death of BMDMs^25^, but in contrast to the reported phosphorylation of RIPK3 and MLKL during PANoptosis^12,13,36^. Instead, our findings indicate that caspase-8 auto-processing is responsible for direct activation of caspase-1 and limits a backup caspase-1 activation pathway that occurs via RIPK3- and MLKL-dependent activation of NLRP3 in the absence of caspase-8 auto-processing.

**Figure 3:**
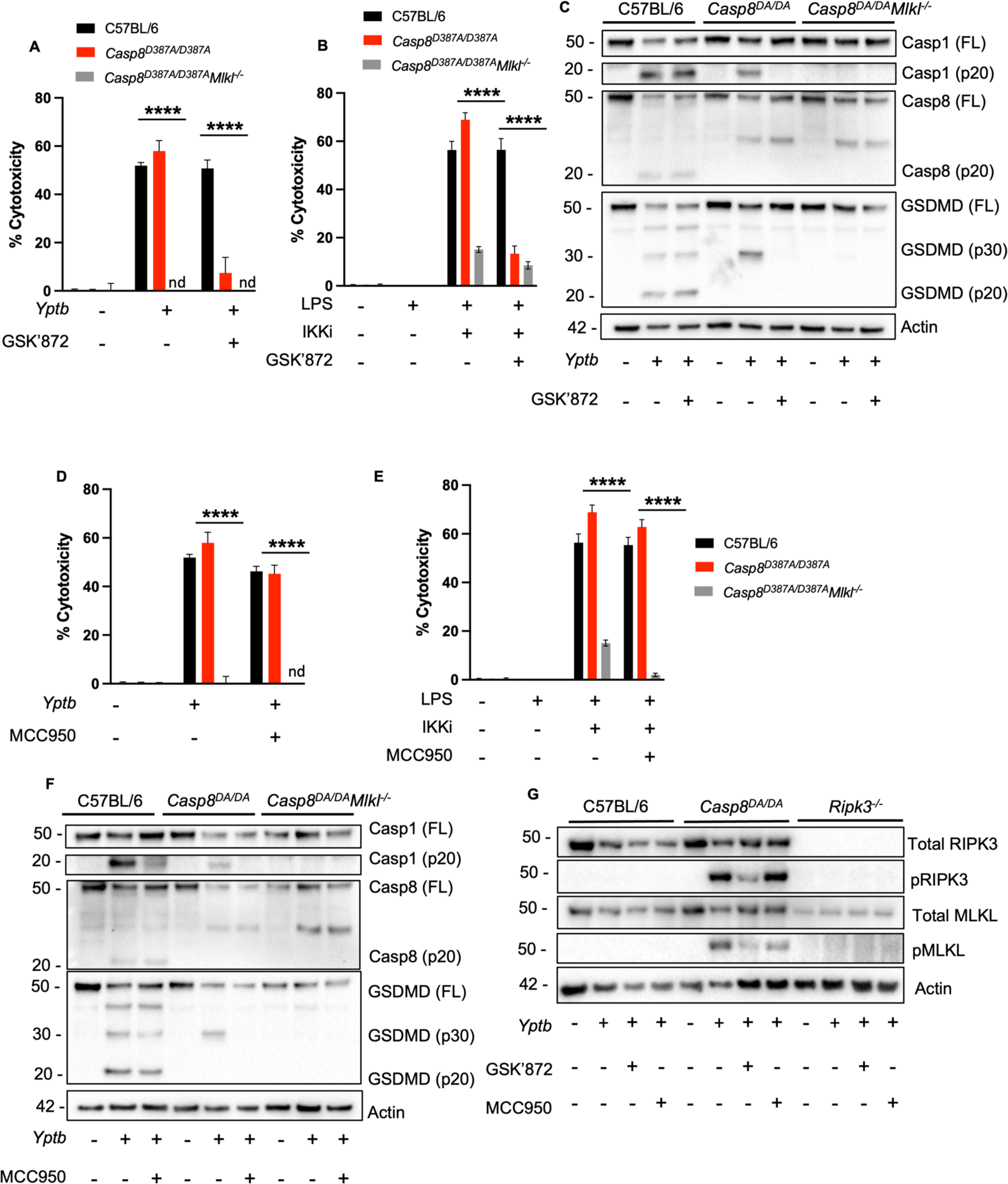
Caspase-8 auto-processing limits RIPK3-mediated necroptosis. (A) C57BL/6, *Casp8^D387A/D387A^*, and *Casp8^D387A/D387A^Mlkl^−/−^* BMDMs were treated with GSK’872 or vehicle control as indicated and infected with WT *Yptb*. Percent cytotoxicity was measured 4 hours post-infection as described. (B) C57BL/6, *Casp8^D387A/D387A^*, and *Casp8^D387A/D387A^Mlkl^−/−^* BMDMs were primed with LPS followed by IKK inhibitor. Prior to IKK inhibitor treatment, BMDMs were treated with GSK’872 or vehicle control. Percent cytotoxicity was measured 5 hours post-infection. (C) Lysates collected 3 hours post-infection were immunoblotted for caspase-1, caspase-8, GSDMD, and β-actin. (D) C57BL/6, *Casp8^D387A/D387A^*, and *Casp8^D387A/D387A^Mlkl^−/−^* BMDMs were treated with MCC950 or vehicle control and were infected with WT *Yptb*. Percent cytotoxicity was measured 4 hours post-infection. (E) C57BL/6, *Casp8^D387A/D387A^*, and *Casp8^D387A/D387A^Mlkl^−/−^* BMDMs were primed with LPS followed by IKK inhibitor. Prior to IKK inhibitor treatment, BMDMs were treated with MCC950 or vehicle control. Percent cytotoxicity was measured 5 hours post-infection. (F) Lysates collected 3 hours post-infection were immunoblotted for caspase-1, caspase-8, GSDMD, and β-actin. (G) C57BL/6, *Casp8^D387A/D387A^*, and *Ripk3^−/−^* BMDMs were treated with MCC950, GSK’872, or vehicle control and were infected with WT *Yptb*. Lysates collected 3 hours post-infection were immunoblotted for total RIPK3, pRIPK3, total MLKL, pMLKL, and β-actin. Nd — not detected, **** p < 0.0001 by two-way ANOVA. Error bars represent the mean +/-SEM of triplicate wells and are representative of three independent experiments.

### ASC speck formation is GSDMD- and NLRP3-dependent and is required for IL-1β processing and release

While our findings indicate that NLRP3 activates caspase-1 downstream of RIPK3/MLKL when caspase-8 activation is disrupted, whether and how NLRP3 might contribute to anti-*Yptb* responses in wild-type BMDMs is unclear. Although the NLRP3 inflammasome is activated in response to *Yersinia* infection or IKK/TAK1 blockade^24,44^, NLRP3 and the adaptor ASC do not contribute to either caspase-8 or caspase-1 activation, GSDMD processing, or cytotoxicity^25^. It has been suggested that co-assembly of ASC, NLRP3, RIPK3, caspase-1 and −8 triggers PANoptosis during *Yersinia* infection^12,13^. However, activation of GSDMD can lead to formation of pores that mediate potassium efflux, a common trigger of the NLRP3 inflammasome that can promote feed-forward activation of caspase-1 downstream of other stimuli^45^. Intriguingly, while we did not observe any differences between WT and *Asc^−/−^* BMDMs in the extent or kinetics of caspase-1, −8, or GSDMD processing, robust caspase-8 processing occurred substantially earlier than processing of caspase-1 or GSDMD (**FIG. 4A**), consistent with a model in which caspase-8 activation occurs upstream of NLRP3-dependent caspase-1 activation. To determine if NLRP3 inflammasome activation occurs downstream of GSDMD pore formation following *Yptb* infection, we assessed NLRP3 activation by the formation of large ASC oligomers that can be visualized via fluorescence microscopy^46^ **(FIG. S2A**). Indeed, transgenic BMDMs expressing ASC-citrine^47^ exhibited robust formation of ASC specks following *Yptb* infection (**FIG. 4B-D, S2B).** The NLRP3-specific inhibitor MCC950 abrogated ASC speck formation (**FIG. 4B, C**), as expected. During *Yersinia* infection, caspase-8 is activated at endosomal membranes by recruitment to RAGulator complexes^48^. Critically, both caspase-8 activation and ASC speck formation were dependent on YopJ activity **(FIG. 4D**). Cytotoxicity remained unchanged in infected WT and *Asc^−/−^* BMDMs even with MCC950 treatment, suggesting that NLRP3 inflammasome activation and ASC speck formation occur downstream of GSDMD pore formation and induction of cell lysis following *Yptb* infection **(FIG. S2C**). To test this hypothesis, we assayed ASC speck formation in *Gsdmd^−/−^* BMDMs following *Yptb* infection **(FIG. 4E, F**). Critically, *Gsdmd^−/−^* BMDMs infected with *Yptb* had a significantly lower frequency of ASC specks relative to wild-type cells in response to *Yptb* infection **(FIG. 4E, F**). Furthermore, MCC950 treatment, or loss of either ASC or GSDMD significantly reduced levels of IL-1β release in response to *Yersinia* infection **(FIG. 4G**). Taken together, our results show that NLRP3 inflammasome activation downstream of caspase-8 and caspase-1-dependent GSDMD pore formation mediates ASC oligomerization and IL-1β release.

**Figure 4:**
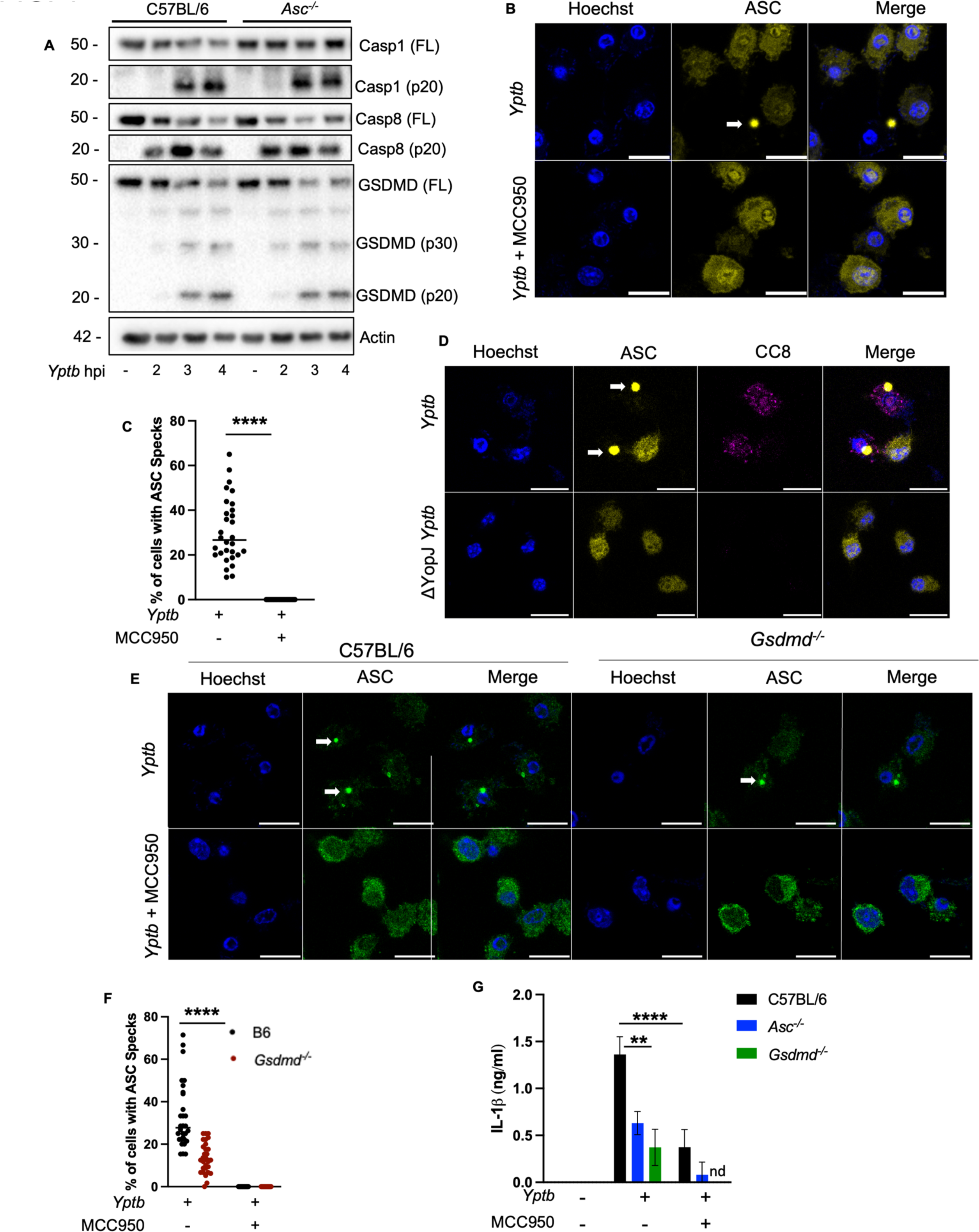
ASC speck formation is GSDMD and NLRP3 dependent and is required for IL-1β processing and release. (A) C57BL/6, and *Asc^−/−^* BMDMs were infected with WT *Yptb*. Lysates collected 3 hours post-infection were immunoblotted for caspase-1, caspase-8, GSDMD, and β-actin. (B) C57BL/6 ASC-citrine BMDMs were treated with MCC950 or vehicle control and were infected with WT *Yptb*. ASC speck formation was analyzed at 4 hours post-infection. (C) Percent of cells with ASC specks was quantified. (D) C57BL/6 ASC-citrine BMDMs were infected with WT and Δ*yopJ Yptb*. Caspase-8 cleavage and ASC speck formation were analyzed at 4 hours post-infection. (E) C57BL/6, and *Gsdmd^−/−^* BMDMs were treated with MCC950 or vehicle control and infected with WT *Yptb*. ASC speck formation was analyzed at 4 hours post-infection via immunofluorescence staining. (F) Percent of cells with ASC specks was quantified. (G) Release of IL-1β into the supernatant was measured by ELISA at 4 hours post-infection in C57BL/6, and *Asc^−/−^*, *Gsdmd^−/−^* BMDMs. ns — not significant, **** p < 0.0001, ** p < 0.001 by two-way ANOVA. Error bars represent the mean +/-SEM of triplicate wells and are representative of three independent experiments.

### Caspase-8 and ASC form separate but functionally linked death complexes

Our findings that ASC speck formation occurs downstream of caspase-8-dependent caspase-1 activation, and that caspase-8 processing precedes caspase-1 processing, suggest that rather than simultaneous engagement of multiple death pathways within a single complex, sequential activation of distinct death complexes occurs during *Yptb* infection. In support, whereas robust caspase-8 activation was detected as early as 2 hours post infection and increased by 4 hours, ASC specks were undetectable at 2h and were only detected at 4 hours post infection (**FIG. 5A-D, S33A**). In addition to activation of caspase-8 and ASC puncta formation being temporally distinct, active caspase-8 and ASC puncta assembly were also spatially distinct, as we observed virtually no colocalization between active caspase-8 puncta and the ASC speck **(FIG. 5C).** Moreover, both caspase-8 activity and ASC speck formation were abrogated upon treatment with the RIPK1 kinase inhibitor Nec-1, whereas the NLRP3 inhibitor MCC950 abrogated ASC speck formation but not active caspase-8 puncta formation **(FIG. 5A, C**). These data indicate that ASC speck formation occurs downstream of caspase-8 activation and is dependent on NLRP3. As both caspase-8 and caspase-1 cleavage of GSDMD can promote NLRP3 activation and ASC speck formation, whether caspase-8 is sufficient, in the absence of caspase-1, to fully activate the NLRP3 inflammasome is unclear. Notably, *Casp1/11^−/−^* ASC-citrine BMDMs exhibited a significant decrease in ASC specks compared to WT ASC-citrine BMDMs in responses to *Yptb* infection, but not in response to LPS/ATP **(FIG. S3B-D**). These data support a model whereby ASC speck formation is upstream of caspase-1 activation in response to LPS/ATP, but downstream of caspase-8-dependent caspase-1 activation in response to YopJ-dependent blockade of immune signaling. Altogether, these data show that during *Yptb* infection, active caspase-8 and ASC complex assembly occur in a kinetically and spatially separable manner.

**Figure 5:**
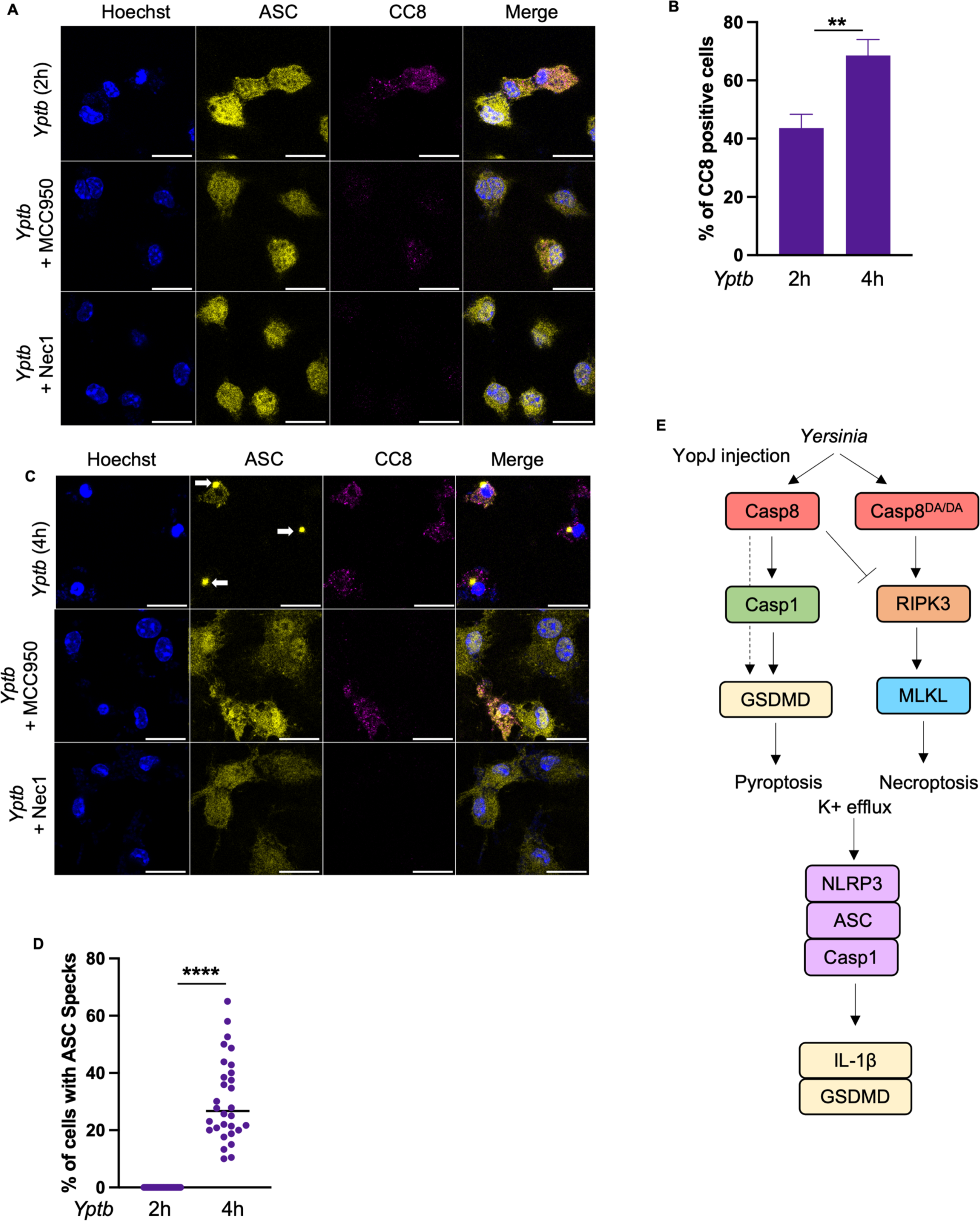
Caspase-8 and ASC form separate but functionally linked death complexes. (A) C57BL/6 ASC-citrine BMDMs were treated with MCC950, Nec-1, or vehicle control and were infected with WT *Yptb*. Caspase-8 cleavage and ASC speck formation were analyzed at 2 hours post-infection. (B) Percent of cleaved caspase-8 positive cells was quantified at 2- and 4 hours post-infection. (C) C57BL/6 ASC-citrine BMDMs were treated with MCC950, Nec-1, or vehicle control and were infected with WT *Yptb*. Caspase-8 cleavage and ASC speck formation were analyzed at 4 hours post-infection. (D) Quantification of percent of cells with ASC specks at 2- and 4 hours post-infection. (E) Graphical representation of findings. ****p < 0.0001 by two-way ANOVA, ** p < 0.05 by unpaired t-test. Error bars represent the mean +/-SEM of triplicate wells and are representative of three independent experiments.

## DISCUSSION

Cell death following blockade of immune signaling kinases TAK1 and IKK by pathogenic *Yersinia* or pharmacological inhibitors is accompanied by activation of both apoptotic and pyroptotic caspases, raising questions about how seemingly distinct forms of cell death can occur simultaneously^11,28^. The activation of pyroptotic and apoptotic caspases, along with the activation of necroptosis when caspase-8 is absent or inhibited, has led to a proposed model in which a unified complex containing regulators of multiple death pathways (pyroptosis, apoptosis, and necroptosis) mediates *Yersinia-* and TAK1 blockade-induced cell death^12,13^. Our findings support an alternative model in which two spatially and temporally distinct, yet functionally linked death complexes assemble in response to *Yptb* infection. Overall our data indicate that caspase-8 initiates downstream responses via direct cleavage of caspase-1, followed by auto-amplification of caspase-1 activation. Caspase-1 activation in response to *Yersinia* infection requires FADD and RIPK1^25^, and the formation of the FADD/RIPK1/caspase-8-containing complex IIa downstream of TAK1 inactivation^49,50^ suggests that caspase-1 activation initially takes place within this complex^51–53^. Caspase-1 and caspase-8 activation within complex IIa also mediates GSDMD cleavage, for which our findings suggest that caspase-1 serves as the primary activator. Our data further demonstrate that GSDMD-dependent activation of the canonical NLRP3-ASC-caspase-1 inflammasome, presumably via potassium efflux, is kinetically and spatially distinct from caspase-8 activation, and is not required for cell death, but is required for secretion of IL-1β. Although caspase-8 can cleave GSDMD to induce pyroptosis in the absence of caspase-1, caspase-8 cannot compensate for lack of caspase-1 or NLRP3 inflammasome activation with respect to IL-1β secretion. Why caspase-1 and the NLRP3 inflammasome are required for IL-1β secretion despite upstream activation of caspase-8 is not clear, but they may enable enhanced or accelerated IL-1β release following *Yptb* infection.

As caspase-1 is cleaved and activated in the absence of inflammasome components during *Yptb* infection^25,49^, we hypothesized that caspase-8 might directly activate caspase-1. Indeed, caspase-8 auto processing and catalytic activity were required for caspase-1 processing in a HEK293 co-expression system. Surprisingly, caspase-1 catalytic activity was also required for its own processing and activation downstream of IKK blockade. This was the case in HEK293 cells as well as in immortalized and primary macrophages from *Casp1^C284A^* mice. Our data suggest that the enzymatic activity of caspase-8 is required to generate a catalytically active scaffold which can then recruit and cleave caspase-1. Our data further indicate that caspase-1 activity is required for its own processing, which likely occurs first within complex IIa, and subsequently within NLRP3 inflammasomes, thereby amplifying the response to enable maximal cleavage of GSDMD, IL-1β, and pyroptosis.

Consistent with prior findings that caspase-1 and −11 are not required for death of BMDMs in response to *Yersinia*^25^, caspase-1 catalytic activity is dispensable for cell death, likely due to caspase-8-dependent cleavage of caspase-3 and −7. GSDME, which is activated by caspase-3 and mediates pyroptosis in other settings^54–56^, also does not contribute to cell death during *Yersinia* infection^29^, indicating that other caspase-3/7 targets are likely responsible. Additionally, consistent with previous reports^23,24^, in the absence of caspase-1, caspase-8-dependent cleavage of GSDMD also occurs and contributes to pyroptosis, although GSDMD cleavage is significantly blunted in the absence of caspase-1.

Simultaneous activation of multiple cell death pathways involving RIPK3 and caspase-8/caspase-1 is proposed to occur during infection by a number of pathogens including *Legionella*, *Francisella,* Influenza, and *Yersinia* infection^11,57–59^. How such a complex assembles remains mysterious, particularly when caspase-8 activity represses RIPK3-dependent necroptosis^9,10^. RIPK3 makes no detectable contribution to *Yersinia-*induced cell death^25,28,60^ and we do not observe any evidence for RIPK3-mediated necroptosis during *Yersinia* infection in the presence of functional caspase-8 **(FIG. 1A, B**). In addition, neither RIPK3 nor MLKL undergo phosphorylation in wild-type cells following *Yersinia* infection or IKK blockade **(FIG. 3G**). Rather, our data favor a model wherein caspase-8 activation and auto-processing downstream of IKK blockade restrains necroptosis, as *Casp8^D387A/D387A^* BMDMs undergo rapid RIPK3 and MLKL phosphorylation, and the cell death that occurs in *Casp8^D387A/D387A^* BMDMs shifts from being RIPK3/MLKL-independent in WT BMDMs to entirely MLKL- and RIPK3 kinase-dependent (**FIG. 3A-C, G**). RIPK3/MLKL-induced programmed necrosis also activates NLRP3, presumably via potassium efflux, thereby providing another route to caspase-1 engagement during *Yersinia* infection, even when caspase-8 cannot be activated (**FIG. 3D-F**). Thus, the coupling of NLRP3 activation to multiple types of lytic pores indicates an important role for backup mechanisms to ensure IL-1β releases and inflammation when immune signaling is inhibited or blocked by pathogen activity.

Our inability to observe RIPK3 and MLKL phosphorylation during *Yersinia* infection of wild-type BMDMs coupled with our observations that caspase-8 activation precedes assembly of ASC specks and detectable caspase-1 activation, suggest that distinct apoptotic and pyroptotic cell death complexes are activated sequentially during *Yersinia* infection. Importantly, caspase-1 is processed in a caspase-8-dependent manner even in *Asc^−/−^* or NLRP3-inhibited BMDMs, indicating that its initial activation takes place within caspase-8-containing complex IIa. Critically, while we observed punctate areas of active caspase-8 throughout the cell following *Yptb* infection, consistent with previous reports^48^, active caspase-8 did not colocalize with ASC specks. Finally, the reduced frequency of ASC specks we observe in the absence of caspase-1 suggests that it serves as the primary activator of GSDMD, which then enables NLRP3 inflammasome activation. Altogether, our study reveals new insight into mechanisms of caspase-8-dependent activation of caspase-1, as well as new understanding of how pyroptotic and apoptotic cell death pathways communicate to mediate anti-microbial host defense.

## MATERIALS AND METHODS

### Cell culture and differentiation of bone marrow-derived macrophages

Bone marrow derived macrophages were isolated and differentiated as previously described^25,49^, in adherence to the NIH Guide for the Care and Use of Laboratory Animals. Briefly, isolated bone marrow cells from 6–10-week-old male and female mice were grown at 37°C, 5% CO2 in 30% macrophage media (30% L929 fibroblast supernatant, complete DMEM). BMDMs were harvested in cold PBS on day 7 and replated in 10% macrophage media onto tissue culture (TC)-treated plates or glass coverslips in TC-treated plates. Transduced iBMDMs from *Casp1^−/−^* mice containing either WT caspase-1, caspase-1 DEAD, or empty vector were previously described and provided by Denise Monack^32^. Primary *Casp1^mlt/mlt^* BMDMs were previously described^35^ and provided by Dr. Olaf Groß. *Casp8^D387A/D387A^Mlkl^−/−^* BMDMs were previously described^43^ and provided by Dr. Doug Green and Dr. Bart Tummers. HEK293T were grown in complete DMEM (supplemented with 10% FBS, 10 mM HEPES, 10 mM Sodium pyruvate, 1% Penicillin/Streptomycin), and maintained in a 37°C incubator with 5% CO2.

### Bacterial culture and *in vitro* infections

Bacterial strains: *Yersinia pseudotuberculosis* (*Yptb*) strain IP2666^61^, *Yptb* ΔYopJ^62^, *Salmonella enterica* serovar Typhimurium strain SL1344 (*S*. Tm)^63^ were all grown as previously described^25^. Briefly, bacteria were grown with aeration and appropriate antibiotics at 28°C (*Yptb*, irgasan) or 37°C (*Salmonella*, streptomycin). *Yptb* strains were induced prior to infection by diluting stationary phase overnight cultures 1:40 in 3 mL of inducing media (2xYT broth, 20 mM Sodium Oxalate, 20 mM MgCl2). Cultures were grown at 28°C for 1 hour and shifted to 37°C for two hours with aeration. *Salmonella* strains were induced prior to infection by diluting the overnight culture 1:40 in 3 mL inducing media (LB broth, 300 mM NaCl), and grown standing for 3 hours at 37°C. Bacterial growth was measured by absorbance at OD600 using a spectrophotometer. Bacteria were pelleted, washed, and resuspended in DMEM or serum-free media for infection. *In vitro* infections were performed at MOI 20 unless otherwise noted. Gentamycin (100 μg/mL) was added one hour post infection for all infections.

### LDH cytotoxicity assay and ELISA

Triplicate wells of BMDMs were seeded in TC-treated 96 well plates. BMDMs were infected with indicated bacterial strains as indicated above. BMDMs were primed with 100 ng/mL LPS for 3 hours followed by 2.5 mM ATP treatment or 5h 10uM IKKi (BMS-345541, Sigma-Aldrich) treatment. BMDMs were primed with 400 ng/mL Pam3CSK4 O/N. BMDMs were treated with 1uM GSK’872 (Invivogen), 1uM MCC950 (Tocris), 60uM Necrostatin-1 (Invivogen) for 30 minutes, 1 hour, and 1 hour prior to infection, respectively. 100 ug/mL gentamycin was added 1 hour post treatment to all infectious experimental conditions. At indicated time points, plates were spun down at 250g, and supernatants were harvested. Supernatants were combined with LDH substrate and buffer (Sigma-Aldrich) according to the manufacturer’s instructions and incubated in the dark for 35 min. Plates were read on a spectrophotometer at 490 nm. Percent cytotoxicity was calculated by background subtraction and normalizing to maximal cell death (1% triton X). To assess IL-1β release, supernatants were diluted 4-fold and applied to Immulon ELISA plates (ImmunoChemistry Technologies) pre-coated with anti-IL-1β capture antibody (eBioscience). Following blocking (1% BSA in 1x PBS), plates were incubated with biotin-linked anti-IL-1β detection antibody (R&D Systems, 1:1000), followed by horseradish peroxidase-conjugated streptavidin. As read-out for IL-1β levels, peroxidase enzymatic activity was determined by exposure to o-phenylenediamine hydrochloride (Sigma) in citric acid buffer. Reactions were stopped with sulfuric acid and absorbance values were read at 490 nm, normalized to mock-transfected cells (negative control).

### HEK293T transfections

Mammalian expression plasmids containing indicated DNA constructs were transfected into HEK293T cells using Lipofectamine 2000 (ThermoFischer) at 1:1 ratio (w/w DNA:Lipofectamine) in Opti-MEM (Gibco). Media was changed to complete DMEM (10% v/v FBS) 6h post-transfection. 24h post-transfection, cells were treated with 1uM AP20187 (dimerizer, ApexBio) in serum-free DMEM for 6h in a humidified incubator at 37°C and 5% CO2 prior to subsequent analysis.

### Western Blotting

BMDMs were seeded in TC-treated 24 well plates (3.0 x10^5^ cells/well). HEK293T cells were seeded in poly-L-lysine-coated TC-treated 24-well plates (2.0 x 10^5^ cells/well) and transiently transfected with appropriate gene constructs as described above. Following infection or treatment in serum-free media, supernatants were harvested, and TCA precipitated overnight at 4°C. Precipitated proteins were pelleted and washed with acetone. The pellet was resuspended in 5X sample buffer (125 mM Tris, 10% SDS, 50% glycerol, 0.06% bromophenol blue, 1% β-mercaptoethanol). BMDMs were lysed in lysis buffer (20 mM HEPES, 150 mM NaCl, 10% glycerol, 1% triton X, 1mM EDTA, pH7.5) plus 1x complete protease inhibitor cocktail and 1x sample buffer (25 mM Tris, 2% SDS, 10% glycerol, 0.012% bromophenol blue, 0.2% β-mercaptoethanol). Lysates and supernatants were boiled and centrifuged at full speed for 5 minutes, were run on 4–12% polyacrylamide gels and transferred to PVDF membrane. Membranes were immunoblotted using the following primary antibodies: β-Actin (Sigma-Aldrich, 1:5000), caspase-1 (gift of Vishva Dixit, Genentech, 1:500), caspase-8 (Enzo, 1:1000), cleaved-caspase-8 (Cell signaling, 1:1000) GSDMD (Abcam, 1:1000), and IL-1β (R&D Systems, 1:1000). Species specific HRP-conjugated secondary antibodies were used for each antibody (1:5000). Membranes were developed using Pierce ECL Plus and SuperSignal West Femto Maximum Sensitivity Substrate (Thermo Fisher Scientific) according to the manufacturer’s instructions. Western blot time-courses were performed in parallel with cytotoxicity assays to accurately interpret protein release before and after overt cell death.

### Fluorescence and confocal microscopy

BMDMs were seeded on circular glass coverslips (Thorlabs, #CG15NH) and allowed to adhere overnight. Cells were then infected or transfected with the indicated DNA constructs (HEK293Ts). At the indicated time points, cells were fixed with 4% PFA for 15 minutes, permeabilized with 0.2% triton X for 10 minutes, and blocked with 5% BSA for 1-2h. BMDMs were stained for cleaved caspase-8 (#8592S Cell signaling, 1:1000) or ASC (#04-147 Millipore, 1:160) overnight at 4°C, Alexa Fluor 647-conjugated anti-rabbit (1:1000), Alexa Fluor 488-conjugated anti-mouse (1:1000) at RT for 1h, and Hoechst at RT for 30 min. Cells were mounted on glass slides with Fluoromount-G (Southern Biotech). Slides were imaged using a Leica SP5-FLIM Inverted confocal microscope with a single z-plane taken per field. Lasers were optimized for GFP (green) Cy5 (far-red), Citrine (yellow), and DAPI (blue). Scale bar = 15 um for all images.

### Image quantification and analysis

Each experiment was conducted in three technical replicates. Within each replicate, 20-30 fields of view were analyzed, with 80-200 cells (BMDMs) per field of view. Specks were defined as distinct high-fluorescence perinuclear clusters of citrine or Alexa Fluor 488 signal. Speck formation frequency was determined as the percentage of citrine-expressing cells that contained one or more specks, using custom macros from ImageJ.

### Statistical analysis

Data were graphed and analyzed using GraphPad Prism. Mean values (± SEM) were compared across triplicate conditions and P values were determined using the appropriate test and are indicated in each figure legend. Studies were conducted without blinding or randomization. Values of *p* < 0.05 were considered statistically significant.

## Acknowledgments

We thank members of the Shin and Brodsky laboratories for helpful scientific discussions. We thank Dr. Denise Monack for generously providing transduced BMDMs from *Casp1^−/−^* mice expressing different caspase-1 mutants, Drs. Doug Green and Bart Tummers for providing *Casp8^D387A/D387A^Mlkl^−/−^* BMDMs, and Dr. Andrew Oberst for providing dimerizable caspase-8 constructs. We thank Leslie King for helpful review and editing of this manuscript. Lastly, we thank Gordon Ruthel at the Penn Vet Imaging Core and for providing helpful insights on fluorescence microscopy.

## Funding

National Institutes of Health grant R01 AI128530 (IEB)

National Institutes of Health grant R01AI139102 (IEB)

National Institutes of Health grant F31 AI172200-01 NRSA (RSW)

Deutsche Forschungsgemeinschaft (DFG, German Research Foundation) SFB 1160 (Project ID 256073931) (BSS, OG)

European Research Council (ERC) Starting Grant 337689 (BSS, OG)

Germany’s Excellence Strategy, through CIBSS - EXC-2189 (Project ID 390939984) (BSS, OG)

## Author contributions

Conceptualization: RSW, IEB

Methodology: RSW, IEB

Investigation: RSW, CKG, BSS

Visualization: RSW, IEB

Funding acquisition: RSW, IEB, PS, BSS, OG

Project administration: IEB

Supervision: IEB, OG, PS

Writing – original draft: RSW, IEB

Writing – review & editing: RSW, IEB, CKG, BSS

## Data and materials availability

All data are available in the main text or the supplementary materials.

## Supplementary Materials

**Figure S1.**
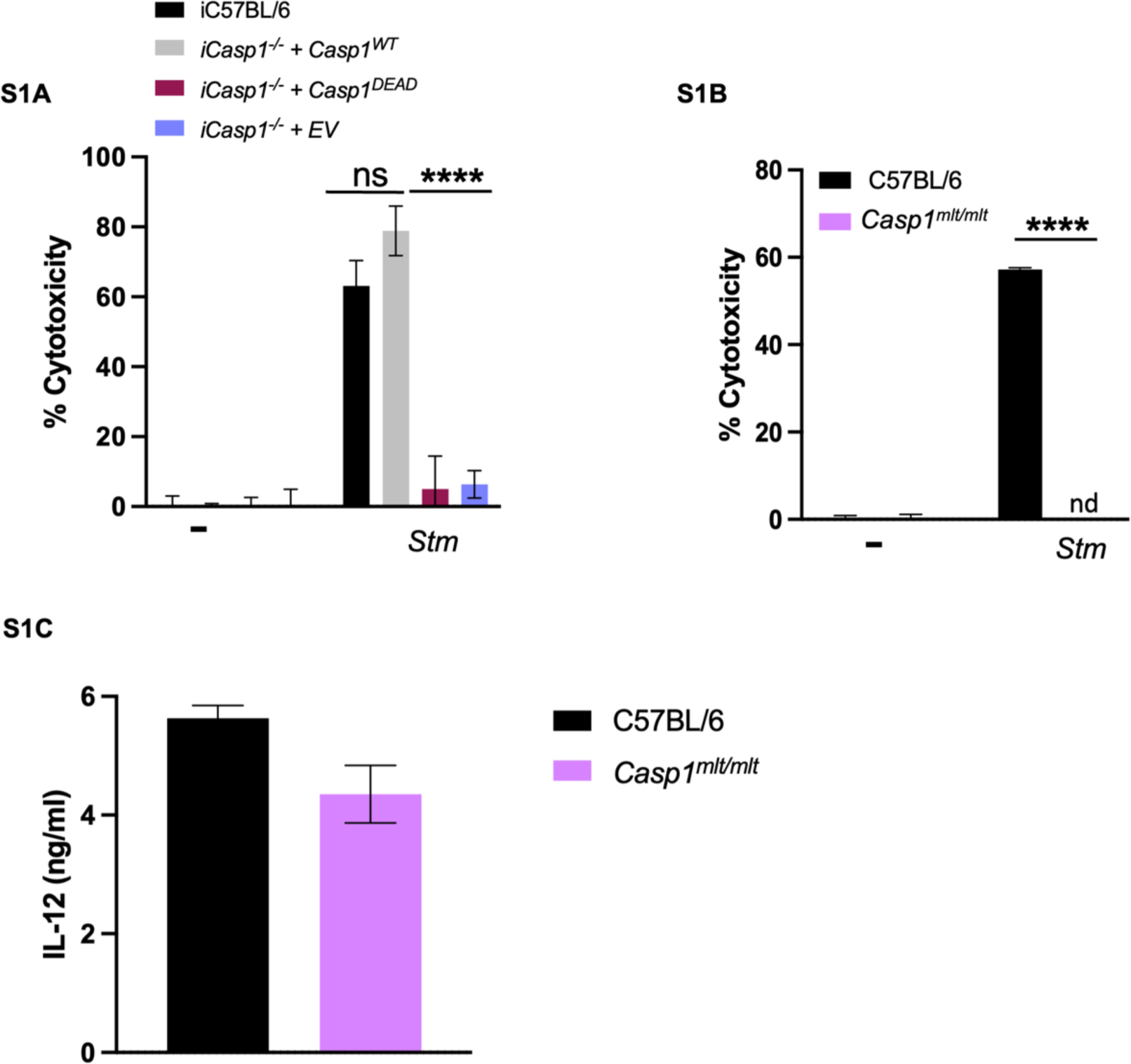
(related to Figure 2)-Caspase-1 catalytic activity is required for cell death during *Stm* infection. (S1A) iC57BL/6, *iCasp1^−/−^ + Casp1^WT^*, *iCasp1^−/−^ + Casp1^DEAD^,* and *iCasp1^−/−^ + EV* iBMDMs were infected with WT *Stm.* Percent cytotoxicity was measured 4 hours post-infection. (S1B) C57BL/6, *Casp1^mlt/mlt^* BMDMs were infected with WT *Stm*. Percent cytotoxicity was measured 4 hours post-infection. (S1C) C57BL/6, *Casp1^mlt/mlt^* BMDMs were infected with WT *Yptb*. Release of IL-12 into the supernatant was measured by ELISA at 4 hours post-infection. ns – not significant. ****p < 0.0001 by two-way ANOVA. Error bars represent the mean +/-SEM of triplicate wells and are representative of three independent experiments.

**Figure S2.**
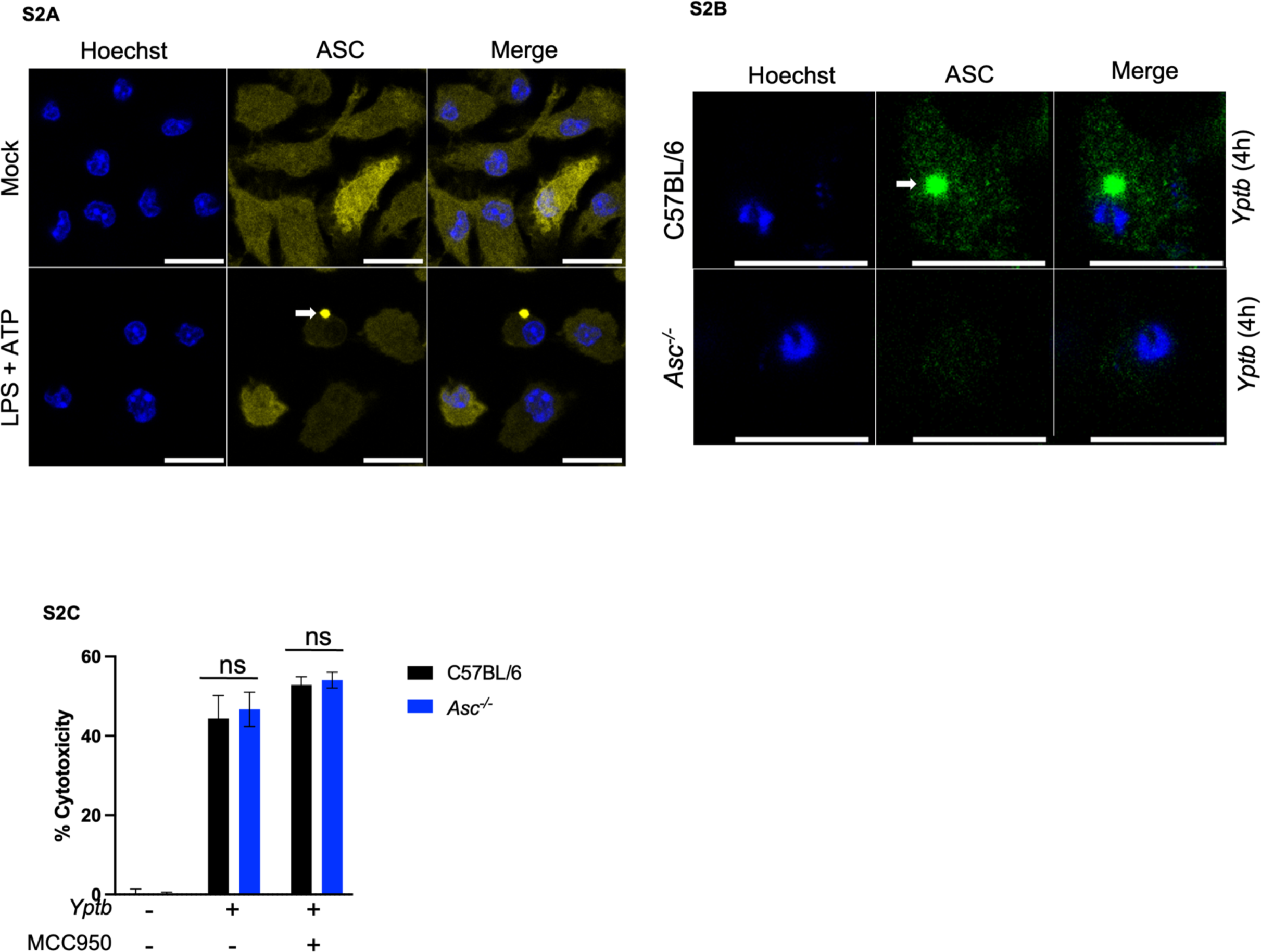
(related to figure 4)-ASC specks can be visualized by fluorescence microscopy. (S2A) C57BL/6 ASC-citrine BMDMs were primed with LPS followed by ATP treatment. ASC speck formation was analyzed 1-hour post ATP treatment. (S2B) C57BL/6, and *Asc^−/−^* BMDMs were infected with WT *Yptb*. ASC speck formation was analyzed 4 hours post-infection. (S2C) C57BL/6, and *Asc^−/−^* BMDMs were treated with MCC950 or vehicle control and were infected with WT *Yptb*. Percent cytotoxicity was measured 4 hours post-infection. ns — not significant. Error bars represent the mean +/-SEM of triplicate wells and are representative of three independent experiments.

**Figure S3.**
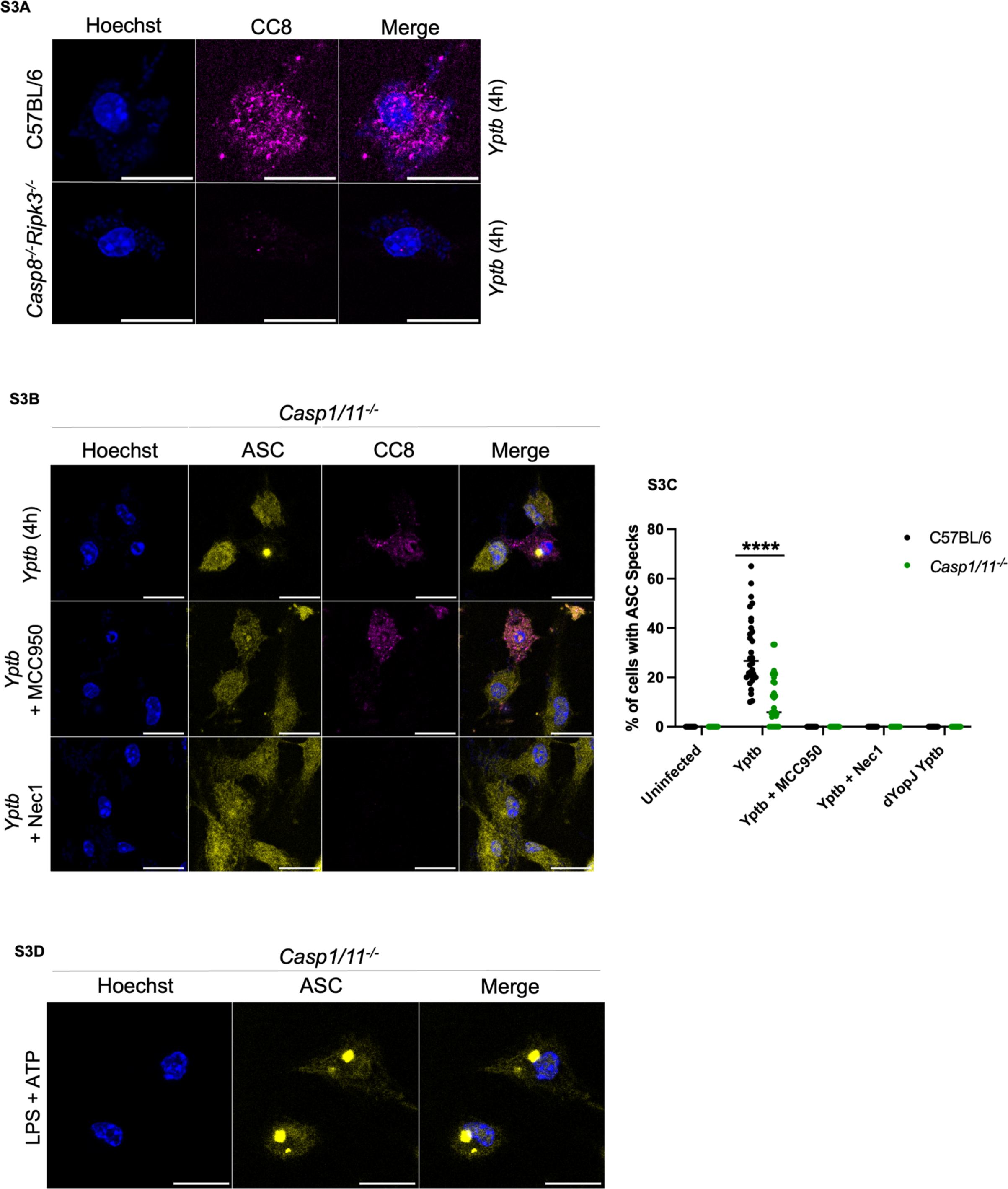
(related to Figure 5)-Absence of casp1/11 results in reduced ASC speck formation. (S3A) C57BL/6, and *Casp8^−/−^Ripk3^−/−^* BMDMs were infected with WT *Yptb*. Caspase-8 cleavage was analyzed 4 hours post-infection. (S3B) 1 hour prior to infection *Casp1/11^−/−^* ASC-citrine BMDMs were treated with MCC950, Nec-1, or vehicle control and were infected with WT *Yptb*. Caspase-8 cleavage and ASC speck formation were analyzed at 4 hours post-infection. (S3C) Quantification of percent of cells with ASC specks for all conditions. (S3D) *Casp1/11^−/−^* C57BL/6 ASC-citrine BMDMs were primed with LPS followed by ATP treatment. ASC speck formation was analyzed 1-hour post ATP treatment. **** p < 0.0001 by two-way ANOVA. Error bars represent the mean +/-SEM of triplicate wells and are representative of three independent experiments.

## References

1. Janeway, C. A. Approaching the asymptote? Evolution and revolution in immunology. Cold Spring Harb. Symp. Quant. Biol. 54 Pt 1, 1–13 (1989).

2. Janeway, C. A. & Medzhitov, R. Innate immune recognition. Annu. Rev. Immunol. 20, 197–216 (2002).

3. Green, D. R. Cell Death: Apoptosis and Other Means to an End, Second Edition. Cold Spring Harbor Laboratory Press (2018).

4. Philip, N. H., Brodsky, I. E., Francis, M., Broz, P. & Anderson, D. Cell death programs in Yersinia immunity and pathogenesis. Front. Cell. Infect. Microbiol. 2, (2012).

5. Nagata, S. Apoptosis and Clearance of Apoptotic Cells. Annu. Rev. Immunol. 36, 489–517 (2018).

6. Cookson, B. T. & Brennan, M. A. Pro-inflammatory programmed cell death. Trends Microbiol. 9, 113–114 (2001).

7. Mariathasan, S. & Monack, D. M. Inflammasome adaptors and sensors: intracellular regulators of infection and inflammation. Nat. Rev. Immunol. 7, 31–40 (2007).

8. Li, J. & Yuan, J. Caspases in apoptosis and beyond. Oncogene 27, 6194–6206 (2008).

9. Kaiser, W. J. et al. RIP3 mediates the embryonic lethality of caspase-8-deficient mice. Nature 471, 368–373 (2011).

10. Oberst, A. et al. Catalytic activity of the caspase-8-FLIP L complex inhibits RIPK3-dependent necrosis. Nature 471, 363–368 (2011).

11. Peterson, L. W. et al. Cell-Extrinsic TNF Collaborates with TRIF Signaling To Promote Yersinia-Induced Apoptosis . J. Immunol. 197, 4110–4117 (2016).

12. Christgen, S. et al. Identification of the PANoptosome: A Molecular Platform Triggering Pyroptosis, Apoptosis, and Necroptosis (PANoptosis). Front. Cell. Infect. Microbiol. 10, 237 (2020).

13. Samir, P., Malireddi, R. K. S. & Kanneganti, T.-D. The PANoptosome: A Deadly Protein Complex Driving Pyroptosis, Apoptosis, and Necroptosis (PANoptosis). Front. Cell. Infect. Microbiol. 0, 238 (2020).

14. Bergsbaken, T., Fink, S. L. & Cookson, B. T. Pyroptosis: Host cell death and inflammation. Nature Reviews Microbiology vol. 7 99–109 (2009).

15. Shi, J. et al. Cleavage of GSDMD by inflammatory caspases determines pyroptotic cell death. Nature 526, 660–665 (2015).

16. He, W. T. et al. Gasdermin D is an executor of pyroptosis and required for interleukin-1β secretion. Cell Res. 25, 1285–1298 (2015).

17. Cornelis, G. R. The Yersinia Yop Virulon, a Bacterial System to Subvert Cells of the Primary Host Defense. Folia Microbiol. (Praha). 43, 253–261 (1998).

18. Cornelis, G. R. & Wolf-Watz, H. The Yersinia Yop virulon: A bacterial system for subverting eukaryotic cells. Molecular Microbiology vol. 23 861–867 (1997).

19. Viboud, G. I. & Bliska, J. B. Yersinia outer proteins: Role in modulation of host cell signaling responses and pathogenesis. Annual Review of Microbiology vol. 59 69–89 (2005).

20. Mukherjee, S. et al. Yersinia YopJ acetylates and inhibits kinase activation by blocking phosphorylation. Science (80-.). 312, 1211–1214 (2006).

21. Palmer, L. E., Hobbie, S., Galá, J. E. & Bliska, J. B. YopJ of Yersinia pseudotuberculosis is required for the inhibition of macrophage TNF-production and downregulation of the MAP kinases p38 and JNK. Mol. Microbiol. 27, 953–965 (1998).

22. Orth, K. et al. Inhibition of the mitogen-activated protein kinase kinase superfamily by a Yersinia effector. Science 285, 1920–3 (1999).

23. Demarco, B. et al. Caspase-8-dependent gasdermin D cleavage promotes antimicrobial defense but confers susceptibility to TNF-induced lethality. Sci. Adv. 6, 3465–3483 (2020).

24. Orning, P. et al. Pathogen blockade of TAK1 triggers caspase-8-dependent cleavage of gasdermin D and cell death. Science (80-.). 362, 1064–1069 (2018).

25. Philip, N. H. et al. Caspase-8 mediates caspase-1 processing and innate immune defense in response to bacterial blockade of NF-B and MAPK signaling. Proc. Natl. Acad. Sci. 111, 7385–7390 (2014).

26. V, S., et al. AIM2 and NLRP3 inflammasomes activate both apoptotic and pyroptotic death pathways via ASC. Cell Death Differ. 20, 1149–1160 (2013).

27. Vajjhala, P. R. et al. The Inflammasome Adaptor ASC Induces Procaspase-8 Death Effector Domain Filaments *. J. Biol. Chem. 290, 29217–29230 (2015).

28. Weng, D. et al. Caspase-8 and RIP kinases regulate bacteria-induced innate immune responses and cell death. Proc. Natl. Acad. Sci. 111, 7391–7396 (2014).

29. Sarhan, J. et al. Caspase-8 induces cleavage of gasdermin D to elicit pyroptosis during Yersinia infection. Proc. Natl. Acad. Sci. 115, (2018).

30. Oberst, A. et al. Inducible dimerization and inducible cleavage reveal a requirement for both processes in caspase-8 activation. J. Biol. Chem. 285, 16632–16642 (2010).

31. Schroder, K. & Tschopp, J. The Inflammasomes. Cell 140, 821–832 (2010).

32. Broz, P., von Moltke, J., Jones, J. W., Vance, R. E. & Monack, D. M. Differential Requirement for Caspase-1 Autoproteolysis in Pathogen-Induced Cell Death and Cytokine Processing | Elsevier Enhanced Reader. Cell Host Microbe 8, 471–483 (2010).

33. Datta, D., McClendon, C. L., Jacobson, M. P. & Wells, J. A. Substrate and Inhibitor-induced Dimerization and Cooperativity in Caspase-1 but Not Caspase-3. J. Biol. Chem. 288, 9971–9981 (2013).

34. Liu, H., Chang, D. W. & Yang, X. Interdimer Processing and Linearity of Procaspase-3 Activation. J. Biol. Chem. 280, 11578–11582 (2005).

35. Schneider, K. S. et al. The Inflammasome Drives GSDMD-Independent Secondary Pyroptosis and IL-1 Release in the Absence of Caspase-1 Protease Activity. Cell Rep. 21, 3846–3859 (2017).

36. Franchi, L., Kanneganti, T.-D., Dubyak, G. R. & Núñez, G. Differential requirement of P2X7 receptor and intracellular K+ for caspase-1 activation induced by intracellular and extracellular bacteria. J. Biol. Chem. 282, 18810–8 (2007).

37. H, W., et al. Mixed lineage kinase domain-like protein MLKL causes necrotic membrane disruption upon phosphorylation by RIP3. Mol. Cell 54, 133–146 (2014).

38. Sun, L. et al. Mixed Lineage Kinase Domain-like Protein Mediates Necrosis Signaling Downstream of RIP3 Kinase. Cell 148, 213–227 (2012).

39. SA, C., et al. Active MLKL triggers the NLRP3 inflammasome in a cell-intrinsic manner. Proc. Natl. Acad. Sci. U. S. A. 114, E961–E969 (2017).

40. KD, G., et al. MLKL Activation Triggers NLRP3-Mediated Processing and Release of IL-1β Independently of Gasdermin-D. J. Immunol. 198, 2156–2164 (2017).

41. Dondelinger, Y. et al. Serine 25 phosphorylation inhibits RIPK1 kinase-dependent cell death in models of infection and inflammation. Nat. Commun. 10, 1729 (2019).

42. Chen, K. W. et al. Extrinsic and intrinsic apoptosis activate pannexin-1 to drive NLRP 3 inflammasome assembly. EMBO J. 38, (2019).

43. Tummers, B. et al. Caspase-8-Dependent Inflammatory Responses Are Controlled by Its Adaptor, FADD, and Necroptosis. Immunity 52, 994–1006.e8 (2020).

44. Y, Z., et al. A Yersinia effector with enhanced inhibitory activity on the NF-κB pathway activates the NLRP3/ASC/caspase-1 inflammasome in macrophages. PLoS Pathog. 7, (2011).

45. Muñoz-Planillo, R. et al. K^+^ efflux is the common trigger of NLRP3 inflammasome activation by bacterial toxins and particulate matter. Immunity 38, 1142–53 (2013).

46. A, S., GL, H., BG, M. & E, L. ASC speck formation as a readout for inflammasome activation. Methods Mol. Biol. 1040, 91–101 (2013).

47. Tzeng, T.-C., et al. A Fluorescent Reporter Mouse for Inflammasome Assembly Demonstrates an Important Role for Cell-Bound and Free ASC Specks during In Vivo Infection. Cell Rep. 16, 571–582 (2016).

48. Zheng, Z. et al. The lysosomal Rag-Ragulator complex licenses RIPK1– and caspase-8– mediated pyroptosis by Yersinia. Science (80-.). 372, (2021).

49. Brodsky, I. E. et al. A Yersinia effector protein promotes virulence by preventing inflammasome recognition of the type III secretion system. Cell Host Microbe 7, 376–387 (2010).

50. Muzio, M. et al. FLICE, A Novel FADD-Homologous ICE/CED-3–like Protease, Is Recruited to the CD95 (Fas/APO-1) Death-Inducing Signaling Complex. Cell 85, 817–827 (1996).

51. Vandenabeele, P., Declercq, W., Van Herreweghe, F. & Vanden Berghe, T. The Role of the Kinases RIP1 and RIP3 in TNF-Induced Necrosis. Sci. Signal. 3, (2010).

52. Feoktistova, M. et al. cIAPs Block Ripoptosome Formation, a RIP1/Caspase-8 Containing Intracellular Cell Death Complex Differentially Regulated by cFLIP Isoforms. Mol. Cell 43, 449–463 (2011).

53. Peterson, L. W. & Brodsky, I. E. To catch a thief: regulated RIPK1 post-translational modifications as a fail-safe system to detect and overcome pathogen subversion of immune signaling. Current Opinion in Microbiology vol. 54 111–118 (2020).

54. Xu, W. feng et al. Gasdermin E-derived caspase-3 inhibitors effectively protect mice from acute hepatic failure. Acta Pharmacol. Sin. 42, 68–76 (2021).

55. Zhang, Z. et al. Caspase-3-mediated GSDME induced Pyroptosis in breast cancer cells through the ROS/JNK signalling pathway. J. Cell. Mol. Med. 25, 8159–8168 (2021).

56. Yu, J. et al. Cleavage of GSDME by caspase-3 determines lobaplatin-induced pyroptosis in colon cancer cells. Cell Death Dis. 10, 193 (2019).

57. Lei, X., Chen, Y., Lien, E. & Fitzgerald, K. A. MLKL-Driven Inflammasome Activation and Caspase-8 Mediate Inflammatory Cell Death in Influenza A Virus Infection. MBio 14, (2023).

58. Gonçalves, A. V. et al. Gasdermin-D and Caspase-7 are the key Caspase-1/8 substrates downstream of the NAIP5/NLRC4 inflammasome required for restriction of Legionella pneumophila. PLOS Pathog. 15, e1007886 (2019).

59. R, P., et al. AIM2/ASC triggers caspase-8-dependent apoptosis in Francisella-infected caspase-1-deficient macrophages. Cell Death Differ. 19, 1709–1721 (2012).

60. Philip, N. H. et al. Activity of Uncleaved Caspase-8 Controls Anti-bacterial Immune Defense and TLR-Induced Cytokine Production Independent of Cell Death. PLoS Pathog. 12, (2016).

61. Zhang, Y., Murtha, J., Roberts, M. A., Siegel, R. M. & Bliska, J. B. Type III Secretion Decreases Bacterial and Host Survival following Phagocytosis of Yersinia pseudotuberculosis by Macrophages. Infect. Immun. 76, 4299–4310 (2008).

62. Lilo, S., Zheng, Y. & Bliska, J. B. Caspase-1 Activation in Macrophages Infected with Yersinia pestis KIM Requires the Type III Secretion System Effector YopJ. Infect. Immun. 76, 3911–3923 (2008).

63. Hoiseth, S. K. & Stocker, B. A. D. Aromatic-dependent Salmonella typhimurium are non-virulent and effective as live vaccines. Nature 291, 238–239 (1981).

